# Genome-wide sequencing uncovers cryptic diversity and mito-nuclear discordance in the *Octopus vulgaris* species complex

**DOI:** 10.1101/573493

**Authors:** Michael D. Amor, Stephen R. Doyle, Mark D. Norman, Alvaro Roura, Nathan E. Hall, Andrew J. Robinson, Tatiana S. Leite, Jan M. Strugnell

## Abstract

Many marine species once considered to be cosmopolitan are now recognised as cryptic species complexes. Mitochondrial markers are ubiquitously used to address phylogeographic questions, and have been used to identify some cryptic species complexes; however, their efficacy in inference of evolutionary processes in the nuclear genome has not been thoroughly investigated. We used double digest restriction site-associated DNA sequencing (ddRADseq) markers to quantify species boundaries in the widely distributed and high value common octopus, *Octopus vulgaris*, comparing genome-wide phylogenetic signal to that obtained from mitochondrial markers. Phylogenetic analyses, genome-wide concordance and species tree estimation based on 604 genome-wide ddRADseq loci revealed six species within the *O. vulgaris* group. Divergence time estimates suggested modern-day species evolved over the last 2.5 ma, during a period of global cooling. Importantly, our study identified significant phylogenetic discordance between mitochondrial and nuclear markers; genome-wide nuclear loci supported *O. vulgaris* sensu stricto and Type III (South Africa) as distinct species, which mtDNA failed to recognise. Our finding of conflicting phylogenetic signal between mitochondrial and nuclear markers has broad implications for many taxa. Improved phylogenetic resolution of *O. vulgaris* has significant implications for appropriate management of the group and will allow greater accuracy in global fisheries catch statistics.

## Introduction

Cryptic diversity is common throughout the animal kingdom, occurring almost homogeneously among metazoan taxa^1^. The introduction of molecular sequencing has greatly improved our understanding of cryptic diversity^2^ and it is clear that DNA barcoding and the use of mitochondrial DNA (mtDNA) markers have been invaluable in revealing many cryptic species complexes^3–5^. However, mtDNA markers have been shown to both underestimate and overestimate species boundaries, particularly among invertebrates^6^. Coalescence estimates based on mtDNA markers are typically poorly correlated with phylogenetic divergence at nuclear genes^7^, and estimates of species diversity based on mtDNA can lead to mischaracterisation of recently diverged species^8^. Although mtDNA barcoding has the potential to accurately delimit species^5^, one study reported a success rate of <30% when using this method^9^. This suggests that mtDNA-based inferences of species-level boundaries among broad ranging taxa may benefit from the inclusion of additional markers. In fact, many authors already recommend that DNA-based species delimitation should employ multiple loci to avoid such biases^8,10,11^.

Several studies have reported discordance between a single or small number of nuclear and mtDNA markers when attempting to delimit cryptic species^12^ (see reference for review). However, few studies have compared species boundaries inferred from genome-wide nuclear markers to those estimated using mitochondrial markers. Genome-wide measures of genetic diversity are becoming increasingly common due to advances in high-throughput Next Generation Sequencing (NGS) technologies, and rapid genotyping of hundreds to thousands of markers from virtually any genome is now possible^13^, even with little or no previously available genetic information^14^. NGS approaches such as restriction site-associated DNA sequencing (RADseq)^15^ are well suited to phylogenetic studies, due to the low cost and high sample throughput relative to other NGS approaches^16–18^. For example, genome-wide data obtained via RADseq have revealed greater phylogenetic resolution of cichlids^19^ and swordtails^20^, when compared to studies using mtDNA markers. This suggests that phylogenetic analyses of a large number of genome-wide markers may be well suited to resolve cryptic species complexes. This is important, as the misidentification of ecologically, medically and/or economically valuable species can have serious consequences for pest control, pathogen resistance, conservation of biodiversity and fisheries management^21^.

*Octopus vulgaris* is a highly-valued fisheries target throughout its range, and its north-west African population constitutes the largest single-species octopus fishery in the world^22^. Based on indistinguishable morphology, *O. vulgaris* was historically considered to have a cosmopolitan distribution, occurring in the Mediterranean Sea and the shallow tropical waters surrounding Australasia, Europe, Africa, Asia and the Americas^23^. However, recent mtDNA-based studies have provided some evidence for the existence of species boundaries among populations of *O. vulgaris*^24–32^. Such mtDNA evidence suggests *O. vulgaris* comprises a complex of morphologically similar but genetically distinct *O. vulgaris*-like species (the ‘*Octopus vulgaris* species complex’), which is composed of several hypothesised species ‘Types’^33^ (Fig. 1). A recent morphology-based investigation of this species complex (and close relatives) supported mtDNA-derived phylogenetic structure^32^. However, no studies of this group have attempted to validate these findings using nuclear data.

**Figure 1.**
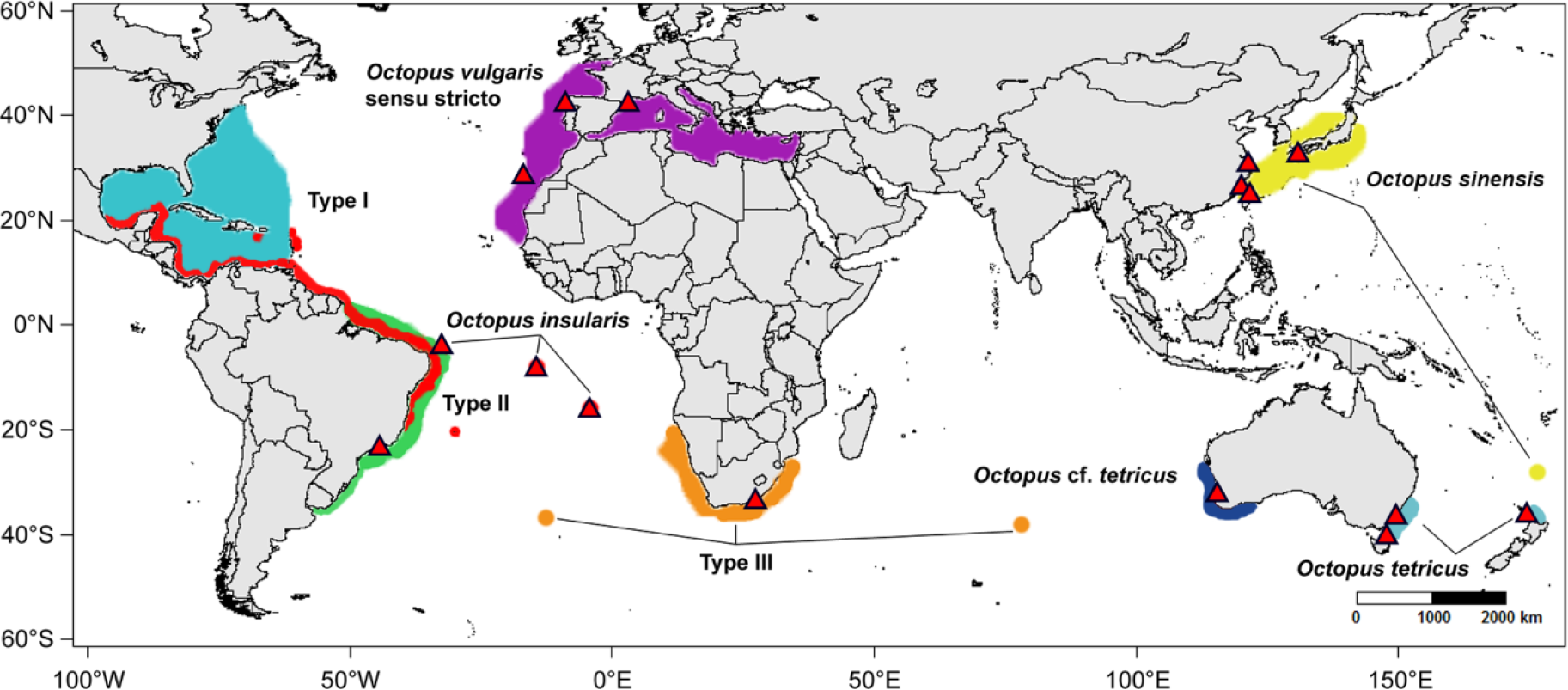
Sampling locations (red triangles) of *Octopus vulgaris* species group [*O. vulgaris* sensu stricto (Mediterranean and eastern North Atlantic), *O. vulgaris* Type II (southern Brazil) *O. vulgaris* Type III (South Africa), *O. tetricus* (east Australia), *O*. cf. *tetricus* (west Australia), *O. sinensis* (Asia)] and *O. insularis* individuals included in the present study. Distributions of *O. vulgaris* sensu stricto and species ‘Types’ are shaded as per Norman *et al*.^33^. Purple, *O. vulgaris* s. s.; blue, Type I (not sampled in our study - Caribbean/ Gulf of Mexico); green, Type II (Brazil); and orange, Type III (South Africa). Distributions of non-*O. vulgaris* species are shaded in yellow: *O. sinensis*, dark blue: *O.* cf. *tetricus*, light blue: *O. tetricus* and red: *O. insularis*. This map was generated using the maps v3.1.1^80^ and mapdata v2.2-6^81^ packages in R v3.1.1^67^.

The poor state of octopus taxonomy is recognised as a major limitation of the global catch statistics reported by the FAO. Of the estimated 100 species of octopus harvested, only four (*O. vulgaris*, *O. maya*, *Eledone cirrhosa* and *E. moschata*) are listed in global statistics^34^. Global production of octopuses exceeds 350,000 tonnes and has a total export value of US$1.07 billion^22^, which surpasses many valuable finfish fisheries. However, the majority of octopus fisheries world-wide are in decline, in part due to overharvesting^34^. Improved taxonomy and identification of cryptic species boundaries is required to improve the reliability of catch estimates and inform sustainable management strategies of this global resource.

Current estimates of diversity within the *O. vulgaris* species complex are likely to underestimate cryptic speciation in this group. In this study, we have employed double digest (dd)RADseq^16^ together with the most extensive global-scale sampling effort to date to investigate the phylogenetic relationships within the *O. vulgaris* species complex and its close relatives. Using genome-wide loci, we have characterised the degree of cryptic diversity within the purportedly widely distributed *O. vulgaris,* and estimated the number of extant species within this group. We contrast our findings with mtDNA-based phylogenies to determine the power of both approaches for resolving phylogenetic relationships. Finally, we use this new phylogenetic information to estimate divergence times among species.

## Results

### Data filtering and assembly

Sequencing of our ddRADseq library resulted in 17.25 million paired-end reads that contained a maximum of a single error per 1,000 bases (Phred quality score of 30). Of the quality paired-end reads, 15.49 million (89.8%) paired reads sufficiently overlapped and therefore were merged into ‘single end data’. Only read one of non-merged data was retained. Unmerged read two sequences were discarded as they were considered to be linked to their associated read one sequence and were lesser quality. Merged and non-merged (read one only) reads were demultiplexed according to their barcodes, which resulted in 13.8 and 1.7 million reads being retained, respectively (90.2% of raw reads). Approximately 9% of the total reads were classified to a microbial database and were discarded; therefore, 91% of the demultiplexed reads were retained. Our shared sequencing run contained 46 samples, 34 were attributed to our study, while the remaining 12 were for an independent project. On average, we obtained 307,316 reads per individual (standard deviation (SD) = 130,692). The 28 *O. vulgaris* group individuals (excluding *O. insularis*) had an average of 336,892 reads per individual (SD = 111,675).

Approximately 69% of our reads mapped to the *O. bimaculoides* genome^35^ across the three datasets (RAD1: all individuals including *O. insularis* (n = 34), RAD2: two to three individuals with highest read number per species (n = 15) and RAD3: only *O. vulgaris group* individuals (n = 26)). The remaining 31% were assembled de novo. This combined approach produced 357,536, 189,678 and 284,924 total putative pre-filtered loci, respectively. Across all three datasets, an average of 3% of the total putative loci were filtered out as they did not meet our set quality criteria and an average of 99% of loci were filtered as they were not present in ≥46% of individuals. The minimum read depth (number of identical reads) per individual of each dataset was 4. Across all loci, the average read depths per individual for the RAD1, RAD2 and RAD3 datasets were 5.9 (SD = 4.4), 4.2 (SD = 3.4) and 5.2 (SD = 4.7), respectively. This resulted in 298 loci with 2,896 informative sites for the RAD1 dataset, 1,060 loci with 7,688 informative sites for the RAD2 data set and 604 loci with 2,976 informative sites for the RAD3 dataset.

### Analysis of mitochondrial DNA derived ddRAD loci

A partial ND2 sequence alignment was obtained after mapping all demultiplexed reads to the mtDNA genome of *O. vulgaris* (Accession: NC_006353.1). Sequence data was obtained for *O. vulgaris* s. s., *O. vulgaris* Type II, *O. vulgaris* Type III, *O. sinensis* and *O. tetricus* which all had a single EcoRI cut site within their mtDNA genome. All *O. insularis* (Brazil, Ascension Island and St Helena) and *O.* cf. *tetricus* individuals were missing data for this region, which may be the result of mutation(s) within the five base pair EcoRI cut site. Maximum Likelihood (ML) analysis of *O. vulgaris* species group individuals placed the five taxa into four distinct monophyletic clades (Bootstrap value [BS] = ≥77; Fig. S1), which is consistent with a previous COI based phylogeny of the group^29^. Each taxon was placed into a distinct clade with the exception of *O. vulgaris* s. s. and *O. vulgaris* Type III which composed a single monophyletic clade.

### Phylogenetic inference using genome-wide ddRAD loci

Maximum likelihood analysis of the *O. vulgaris* group and *O. insularis* individuals using the RAD2 data set (1060 loci) supported seven phylogenetic clades (Fig. 2). *Octopus insularis* formed a highly supported monophyletic clade separated from the *O. vulgaris* group (BS = 100) by a large branch length. *Octopus vulgaris* Type II (Brazil) was placed as sister taxon to the remaining *O. vulgaris* group members and was therefore selected to root the phylogeny for subsequent analyses including only *O. vulgaris* group individuals.

**Figure 2.**
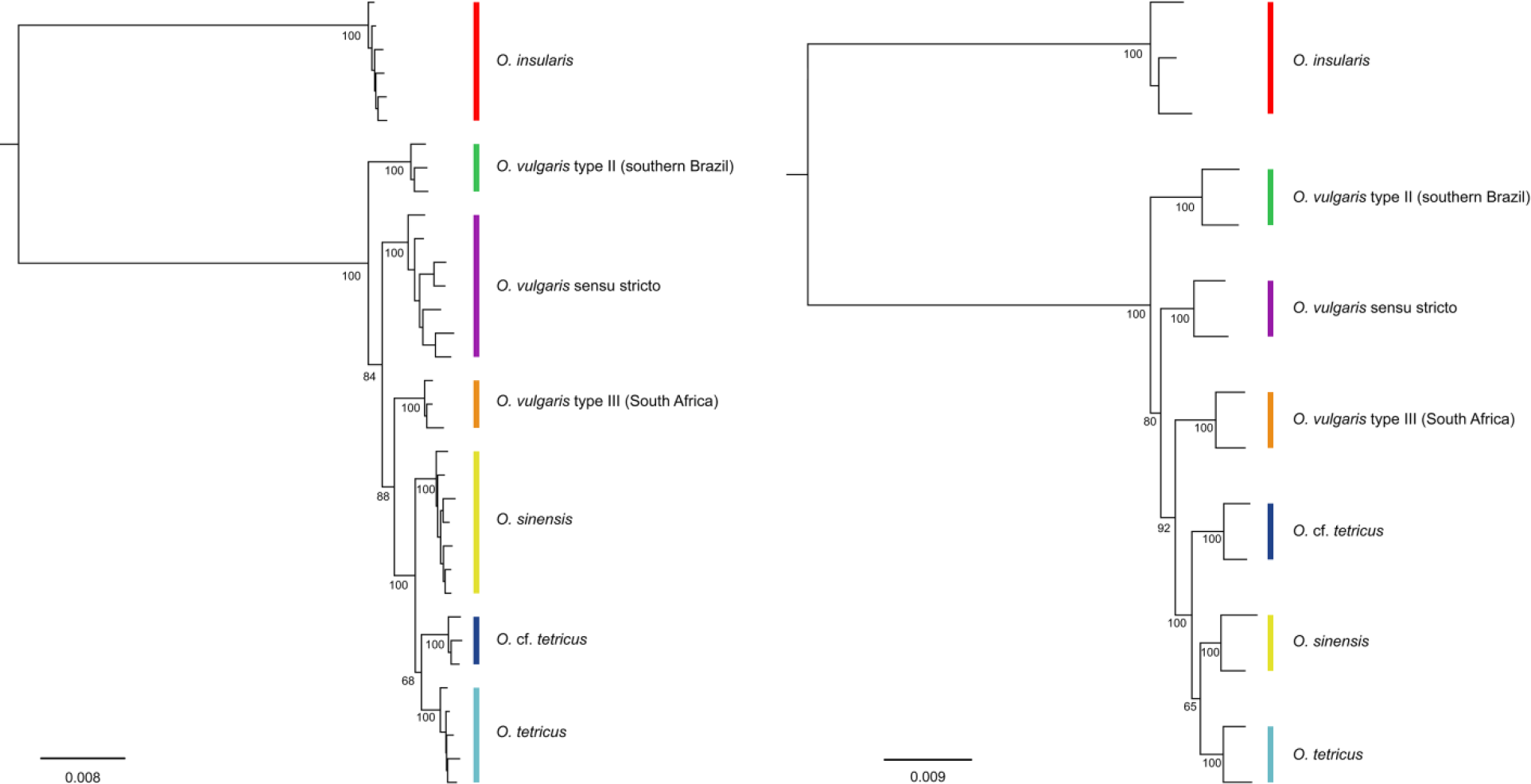
Maximum likelihood phylogenies depicting the relationships among members of the *Octopus vulgaris* species complex, *O. insularis*, *O. tetricus* and *O.* cf. *tetricus*. The phylogeny on the left is based on all 34 individuals sequenced and included 298 loci (RAD1 dataset). The phylogeny on the right is based on the 15 individuals, with 2-3 representatives with the highest number of reads sequenced from each species included. This phylogeny included 1060 loci (RAD02 dataset). Both analyses were conducted using the GTR+G evolutionary model in RAxML v8.0.19 (Stamatakis 2014). ML bootstrap values are displayed below major nodes. Differences among topologies include support at major nodes and the placement/relationships among clades including individuals representing *O. sinensis*, *O. tetricus* and *O.* cf. *tetricus* (although these clades are supported as being distinct (BS = 100). Colours correspond to Fig. 1.

Maximum likelihood analysis of the *O. vulgaris* group individuals (excluding *O. insularis*) based on the RAD3 data set (604 loci) resulted in six highly supported monophyletic clades (Fig. 3). Five of the six phylogenetic clades corresponded to the species previously identified based on differences in morphology^32^. These included *O. vulgaris* Type II (BS = 100), *O. vulgaris* s. s. (BS = 100), *O. sinensis* (BS = 100), *O. tetricus* (BS = 100) and *O.* cf. *tetricus* (BS = 100). Furthermore, individuals from South Africa (*O. vulgaris* Type III) also formed a distinct monophyletic clade (BS = 100).

**Figure 3.**
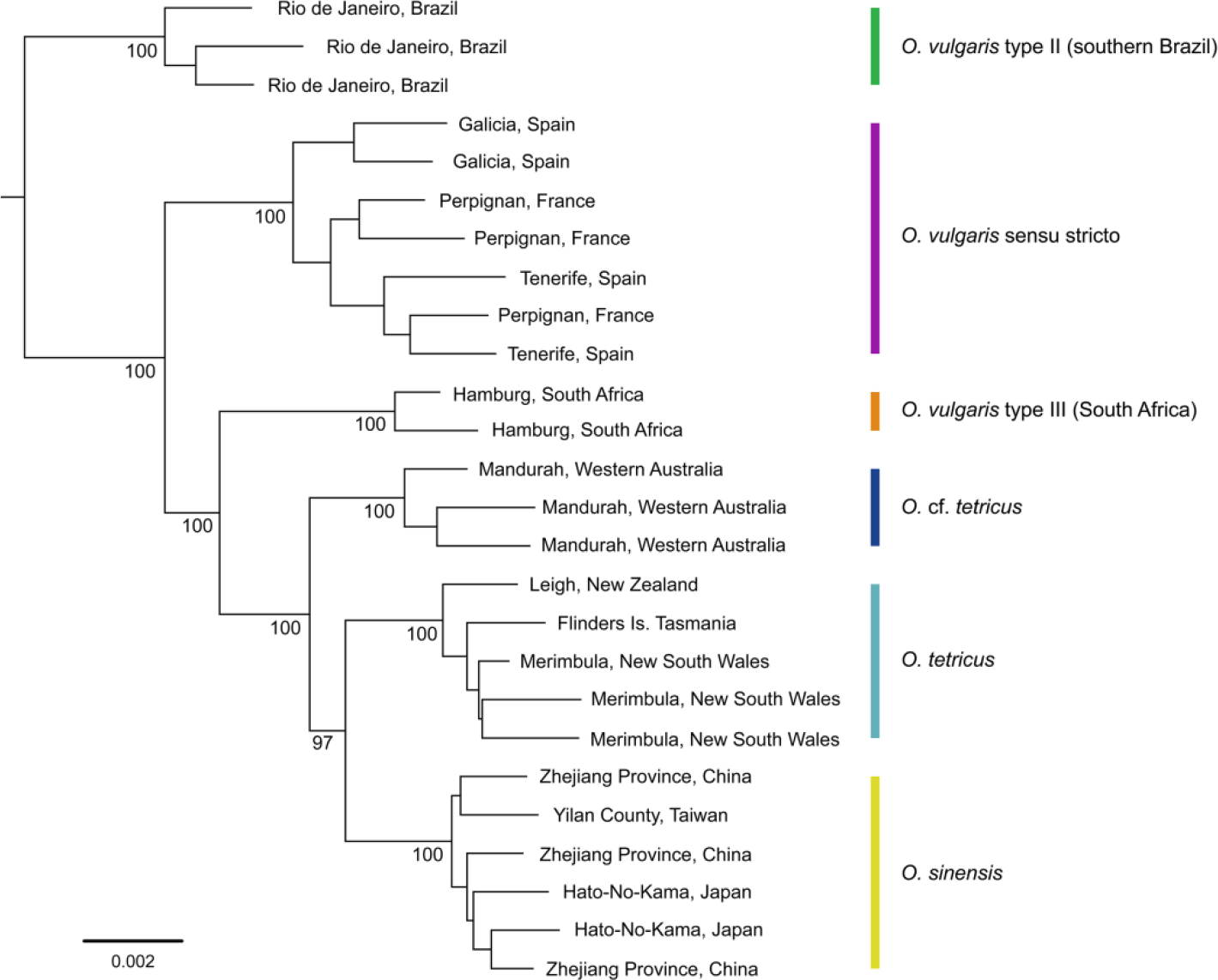
Maximum likelihood phylogeny depicting the relationships among members of the *Octopus vulgaris* species group. Analysis was conducted using 604 concatenated double digest RADseq loci using the GTR+G evolutionary model in RAxML v8.0.19^57^. ML bootstrap values are displayed below major nodes. Tip labels correspond to sampling region in Table 2. Colours correspond to Fig. 1.

### Phylogenetic concordance among genome-wide loci and hypothesis testing of alternate topologies

To investigate the support for sub-optimal trees among the 604 loci from the RAD3 data set, we performed ‘Partitioned RAD’ analysis^36^ using a nearest-neighbour interchange (NNI) approach. The RAxML best tree was shown to be the most reliable representation of the RAD3 data-matrix as it was favoured by the greatest number of loci, and disfavoured by fewer than 10 loci. Our phylogeny (Fig. 3) contrasted previous mtDNA-based topologies, therefore, we performed topological comparisons via an Approximately Unbiased (AU) test, which showed that our RAxML best tree was the highest ranked topology (Posterior Probability [PP] = 1; Table 1). The first constrained topology forced *O. sinensis* to be sister taxon to *O. tetricus* and *O.* cf. *tetricus*, as previously reported^30^. This tree was ranked second and could not be rejected as a significantly worse topology (p = 0.051), however, the support for this tree was far lower compared to the first ranked tree (PP = <0.001). The tree constraining the monophyly of *O. vulgaris* Type III and *O. vulgaris* s. s. (based on mtDNA analyses) was ranked third. This tree was significantly less likely (p = 0.004) than our optimal topology, and the probability supporting it was substantially lower (PP = <0.001). Forcing the monophyly of *O. vulgaris* Type III and *O. vulgaris* s. s. resulted in a sister taxon relationship of two distinct clades, and did not result in the same single clade, ‘conspecific’ relationship obtained via mtDNA-based analyses.

**Table 1.** Approximate Unbiased (AU) test results comparing our phylogenetic topology (Fig. 3) with contrasting results from previous studies. The Bayesian Posterior Probability (PP) calculated under the Bayesian Information Criterion (BIC) is also shown. Topologies are rejected with a p-value of <0.05.

| Tree rank | Constraint | p-value | PP | Reference |
| --- | --- | --- | --- | --- |
| 1 | No constraint | 0.951 | 1 |  |
| 2 | *O. sinensis* as sister taxon to *O. tetricus* and *O. cf. tetricus* | 0.051 | 1E-025 | Amor *et al.*<sup>30</sup> |
| 3 | Forced monophyly of *O. vulgaris* s. s. and *O. vulgaris* Type III | 0.004 | 1E-045 | Amor *et al.*<sup>29</sup> |

**Table 2.**
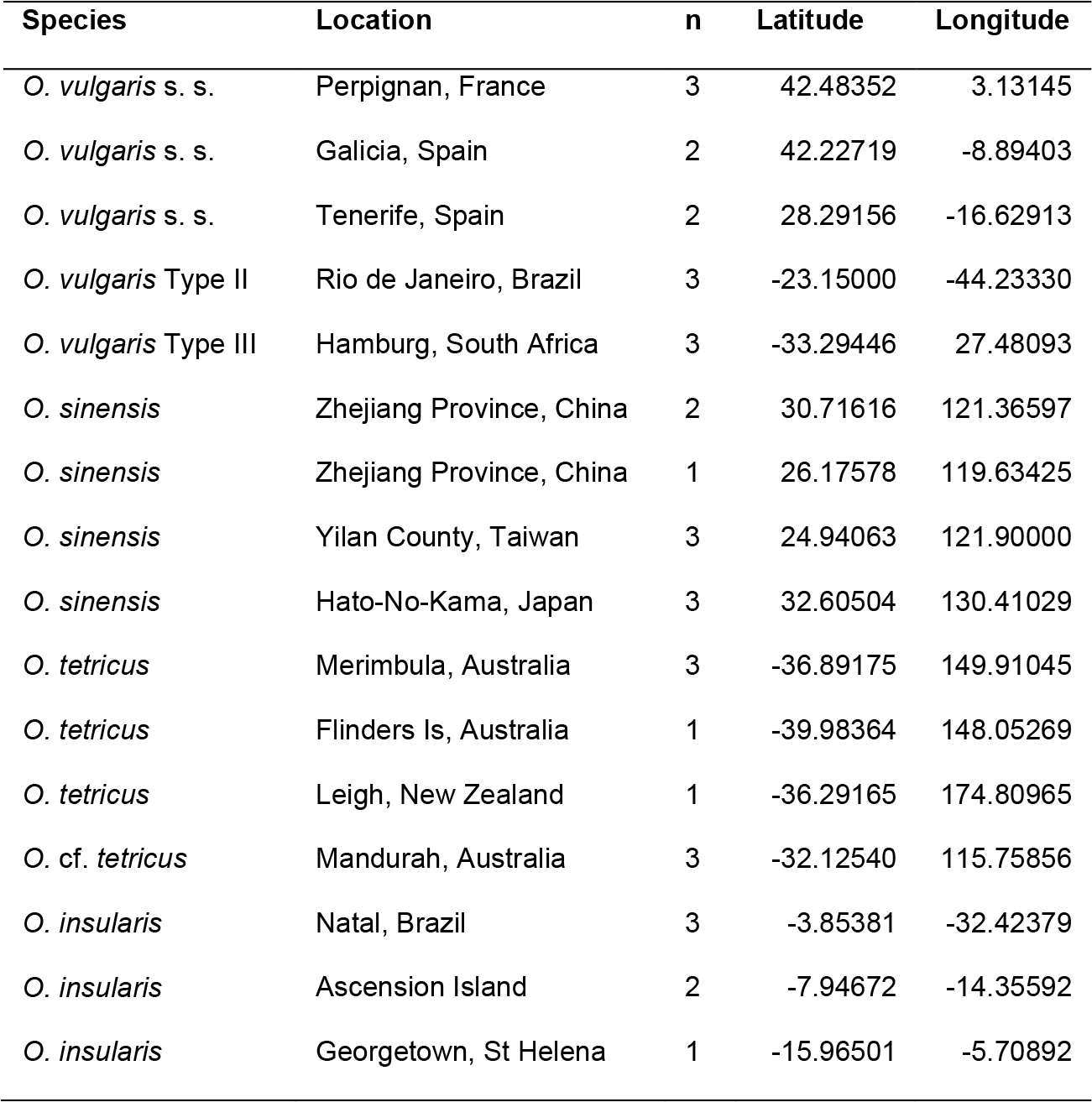
Sampling location of *Octopus vulgaris* species group, and *O. insularis* individuals included in the present study.

| Species | Location | n | Latitude | Longitude |
| --- | --- | --- | --- | --- |
| *O. vulgaris* s. s. | Perpignan, France | 3 | 42.48352 | 3.13145 |
| *O. vulgaris* s. s. | Galicia, Spain | 2 | 42.22719 | -8.89403 |
| *O. vulgaris* s. s. | Tenerife, Spain | 2 | 28.29156 | -16.62913 |
| *O. vulgaris* Type II | Rio de Janeiro, Brazil | 3 | -23.15000 | -44.23330 |
| *O. vulgaris* Type III | Hamburg, South Africa | 3 | -33.29446 | 27.48093 |
| *O. sinensis* | Zhejiang Province, China | 2 | 30.71616 | 121.36597 |
| *O. sinensis* | Zhejiang Province, China | 1 | 26.17578 | 119.63425 |
| *O. sinensis* | Yilan County, Taiwan | 3 | 24.94063 | 121.90000 |
| *O. sinensis* | Hato-No-Kama, Japan | 3 | 32.60504 | 130.41029 |
| *O. tetricus* | Merimbula, Australia | 3 | -36.89175 | 149.91045 |
| *O. tetricus* | Flinders Is, Australia | 1 | -39.98364 | 148.05269 |
| *O. tetricus* | Leigh, New Zealand | 1 | -36.29165 | 174.80965 |
| *O. cf. tetricus* | Mandurah, Australia | 3 | -32.12540 | 115.75856 |
| *O. insularis* | Natal, Brazil | 3 | -3.85381 | -32.42379 |
| *O. insularis* | Ascension Island | 2 | -7.94672 | -14.35592 |
| *O. insularis* | Georgetown, St Helena | 1 | -15.96501 | -5.70892 |

### Species tree estimation

Species tree estimation supported the presence of six species within the *O. vulgaris* group (Fig. 4). Each species was highly supported (BS = >99.1), with *O. vulgaris* s. s.,*O. vulgaris* Type II, *O. vulgaris* Type III, *O.* cf. *tetricus*, *O. tetricus* and *O. sinensis* supported as distinct species.

**Figure 4.**
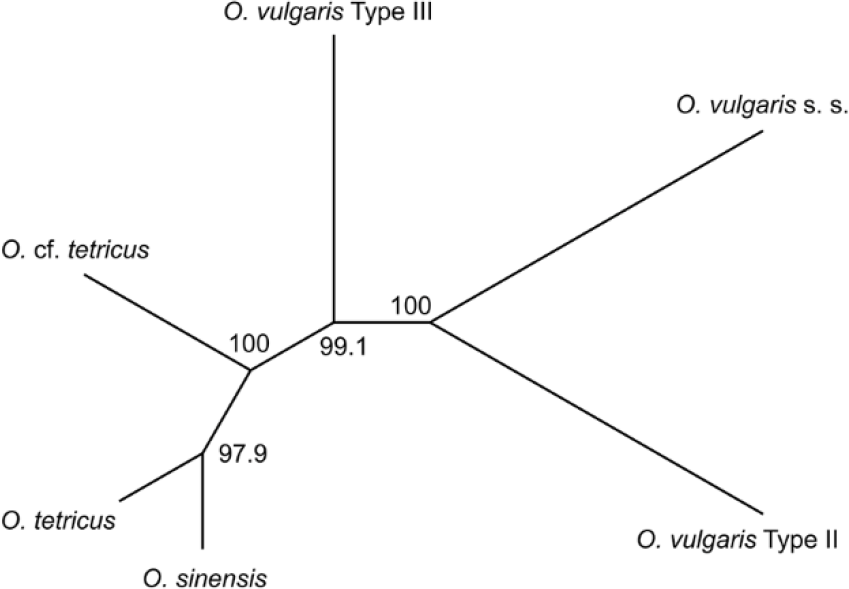
Species tree estimation supporting six species within the *Octopus vulgaris* species group. Analysis was based on 604 concatenated RAD loci using the *SVDQuartets* algorithm^69^ implemented in *PAUP v4.0a146 for Unix/Linux*^70^.

### Divergence time estimation

Divergence times were estimated among the six species identified in the present study; *O. vulgaris* s. s., *O. vulgaris* Type II (Brazil), *O. vulgaris* Type III (South Africa) *O. tetricus*, *O.* cf. *tetricus* and *O. sinensis*. The *O. vulgaris* group was estimated to have evolved within the past 2.5 million years, throughout the Pleistocene (Fig. 5).

**Figure 5.**
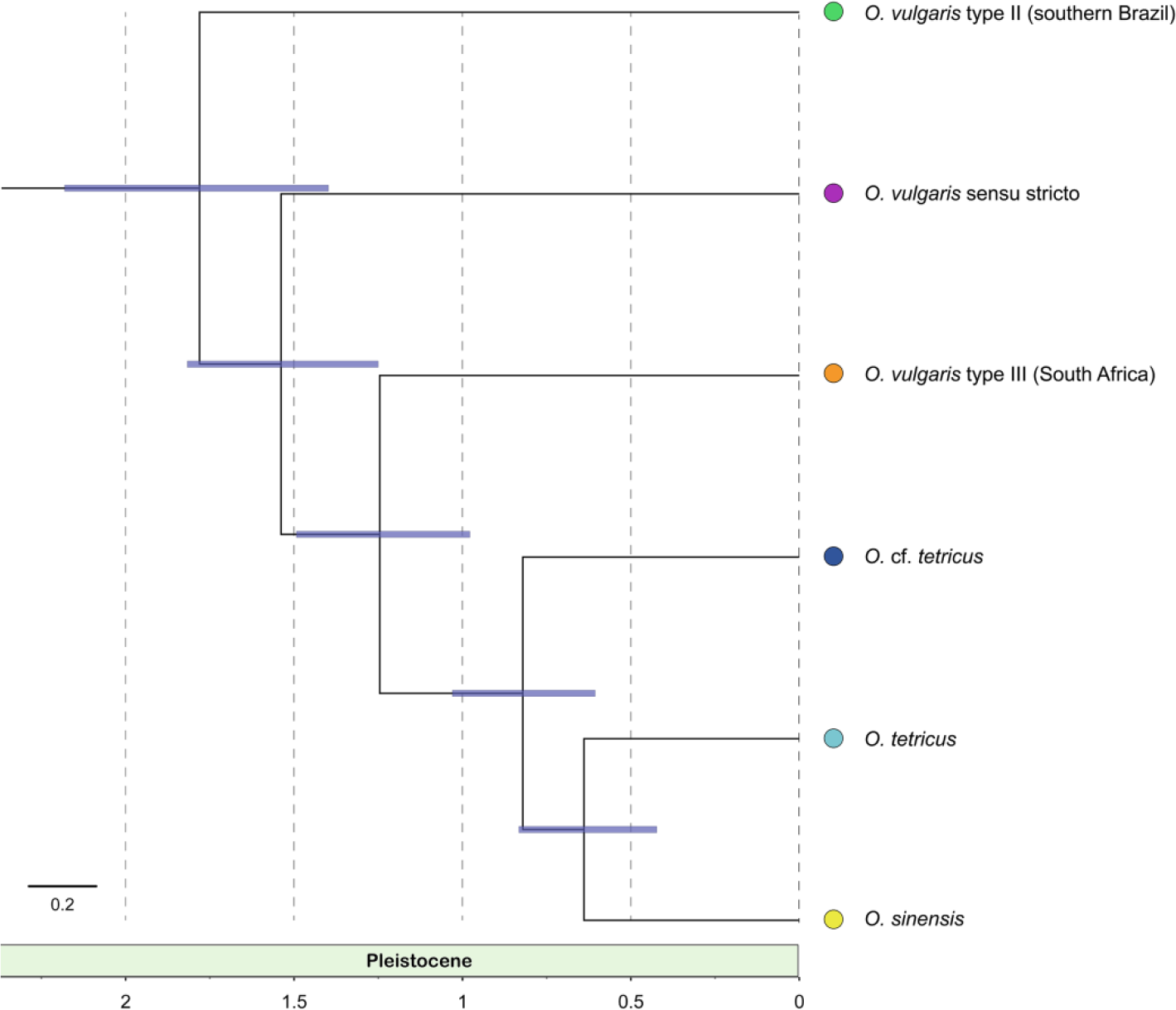
Divergence time estimation within the *Octopus vulgaris* group. The topology presented is based on combining three independent runs that were all calculated using the rate of evolution for cephalopods (0.0036 neutral substitutions per million years). Node bars represent 95% highest posterior densities for divergence estimates and estimate that the *O. vulgaris* species group evolved within the last 2.5 million years. Colours correspond to Fig. 1.

## Discussion

Here, we demonstrate the successful application of ddRADseq for resolving phylogenetic and species-level questions among cryptic taxa. Despite lacking an available reference genome, this approach enabled us to sample hundreds of loci in a random, yet repeatable, manner from across the genome of each individual. Our resulting data-matrix is the most comprehensive assembled so far to investigate species diversity within the *O. vulgaris* species complex. In comparison to previous attempts, which relied solely on mtDNA, our analyses yielded greater phylogenetic resolution and higher clade support. We recommend future studies investigating cryptic diversity use genome-wide approaches such as ddRADseq because of these advantages.

We analysed genome-wide RAD loci to investigate the phylogenetic and species-level relationships within the highest value fisheries target species of octopus in the world. Our study provides evidence for the existence of three cryptic species within the *O. vulgaris* species complex, which has traditionally been considered a single taxonomic unit; *O. vulgaris* s. s. (Mediterranean and north-east Atlantic), *O. vulgaris* Type II (southern Brazil) and *O. vulgaris* Type III (South Africa). We also support that *O. insularis* is distinct from the *O. vulgaris* species group, as previously published^26^. In accordance with previous studies, phylogenetic relationships and species tree estimation in our study support the distinct species status of *O. tetricus*, *O.* cf. *tetricus* and *O. sinensis*, and confirm that New Zealand’s *O. gibbsi* is a junior synonym of east Australia’s *O. tetricus*^30^.

Our findings based on nuclear DNA provide greater resolution than previous studies using mtDNA alone to investigate phylogenetic relationships within the *O. vulgaris* group. Although these previous studies have shown some phylogenetic structure among species^24,25,28–31^, our finding of clear species-level differentiation between *O. vulgaris* s. s. and *O. vulgaris* Type III is a novel discovery. The recent morphological investigation of this species group^32^ did not include *O. vulgaris* Type III individuals. Our study provides evidence that this taxon is a distinct species and, therefore, a comparison of its morphology with other members of the *O. vulgaris* group is required for taxonomic purposes.

Based on analyses of mtDNA, *O. sinensis* has the widest distribution of any taxa within the *O. vulgaris* group. A previous analysis of mtDNA suggests individuals from Asia and the Kermadec Islands are conspecifics^31^, despite allopatric distributions in the North and South Pacific Ocean. Preliminary mtDNA-based analyses also suggest that individuals from the Caribbean/Gulf of Mexico (Type I) and South Brazil (Type II) are conspecifics^37^, which also appears to be true for *O. insularis* from the Caribbean and eastern North Brazil^38,39^. Both of the above examples suggest these taxa can maintain gene-flow despite connectivity requiring trans-tropical dispersal. Since we did not include samples from the Caribbean/Gulf of Mexico and Kermadec Islands in our study, and given that our analyses find that mtDNA underestimates cryptic species diversity in this species complex, these findings require further validation and complimentary morphological analyses.

Biogeographic inconsistencies between mtDNA and nuclear DNA typically rule out lineage sorting as a mechanism contributing to discordance^12^. If nuclear DNA shows greater levels of geographic structure than mtDNA analyses, this discordance is often the result of isolation followed by secondary contact and hybridisation. We therefore suggest that the absence of any mtDNA-based phylogenetic signal between *O. vulgaris* s. s. and South African *O. vulgaris* Type III may be the result of hybridisation following secondary contact. Previously, two distinct mtDNA lineages of *O. vulgaris* Type III were identified along the South African coastline^40^. The dominant lineage was abundant along the entire South African coastline, whilst the rarer lineage was reported in two individuals from the south-eastern extreme of the known distribution (Durban). If our hypothesis is accurate, there may be a selective advantage for the foreign (*O. vulgaris* s. s.) mtDNA throughout South Africa as it appears to be either at, or near, fixation.

The relationships we identified among *O. sinensis*, *O. tetricus* and *O.* cf. *tetricus* using nuclear DNA contrasts previous mtDNA-based analyses. Analyses of Amor *et al.*^30^ showed *O. sinensis* (treated as Asian *O. vulgaris*) was sister taxon to a clade composed of *O. tetricus* and *O.* cf. *tetricus*. Our study instead suggests that *O.* cf. *tetricus* is sister taxon to a clade containing *O. tetricus* and *O. sinensis*, since this topology was significantly more probable. Given that our results are based on genome-wide evidence, we consider our current findings to be a more reliable representation of the true relationships among these taxa.

Our phylogenetic reconstruction also showed overall improvements in support values throughout the tree when compared to previous mtDNA-based topologies. In particular, previous mtDNA-based studies found poor support for the clade containing *O. vulgaris* Type II individuals^29,41^. Datasets with multiple unlinked nuclear DNA loci have a greater ability to resolve phylogenetic relationships in comparison to mtDNA alone^10,11^. Our findings highlight the advantages of using multiple genome-wide loci when investigating species-level relationships in the *O. vulgaris* group, with important implications for future investigations of cryptic diversity among other taxa. Despite these advantages for resolving phylogenetic relationships among closely related taxa, a short-coming of RADseq studies is that an increase in phylogenetic distance also increases locus loss due to mutations at restriction sites^18,36^. However, complimenting ddRADseq with target capture sequencing has shown promise for addressing allelic dropout^42^.

Throughout the evolutionary history of the *O. vulgaris* group, the global geography was relatively similar to the present, however there have been several changes to the global climate^43^. Our estimates of divergence times suggest that modern day *O. vulgaris* group taxa evolved from their MRCA during the last 3 Ma. Over this time, the climate has changed from relatively warm with a reduced equatorial-polar temperature gradient to relatively cool with an increase in Antarctic ice volume and the establishment of the Arctic ice sheet^43,44^. Present day *O. vulgaris* taxa have sub-tropical to temperate distributions in both hemispheres, or inhabit the deeper and cooler water in tropical regions^40,45^. Our study suggests these taxa arose from a common ancestor that inhabited shallow equatorial waters. Our estimates of divergence time suggest modern anti-tropical distributions originated during a period of global cooling, rather than the poleward range shift of each species being the result of increasingly warm and less suitable equatorial temperatures. Anti-tropical divergence in this group may be explained in part by the weakening of ocean currents at this period^46^, which would likely have restricted gene-flow of previously connected populations by limiting larval dispersal via ocean currents.

Our divergence time estimates predict that the *O. vulgaris* group evolved during the last 2.5 Ma. This time period follows warming of global oceans, where temperatures increased between 2-3°C at mid-latitudes and 5-10°C in higher latitudes from 6-3.2 Ma^46,47^. Such temperature increases may have created unsuitably warm climates at low to mid-latitudes, potentially driving a poleward range shift and the isolation of northern and southern hemisphere populations. A poleward shift in distribution may have impeded the transport of larvae between continents via strong equatorial currents. Oceanic gyres in the Indian, southern Atlantic and northern Atlantic Oceans transport larvae among continents, but are strong barriers to planktonic dispersal between hemispheres^48^. Despite the planktonic larvae of *O. vulgaris* s. s. being primarily oceanic^49^, our phylogenetic results support previous studies that suggest *O. vulgaris* group taxa can only maintain gene-flow across distances of 2000-3000 km^29,30^, and appear unable to maintain gene-flow among major continents. *Octopus insularis* inhabits shallow tropical waters throughout the West Atlantic (Espirito Santo state (southern Brazil) to the Gulf of Mexico^37,38,39^ and mid-Atlantic oceanic islands^26,29,50^. As this tropical species is a distinct phylogenetic lineage from the *O. vulgaris* group, how the above mentioned historical temperature increases and fluctuation in current strength influenced its distribution requires future attention.

We have provided compelling evidence for cryptic speciation within the *O. vulgaris* species complex, which has increased our understanding of the geographic boundaries of this species complex and its close relatives. Our study has revealed that octopuses currently being treated as a single *O. vulgaris* species actually represent three distinct species (*O. vulgaris* s. s., *O. vulgaris* Type II and *O. vulgaris* Type III). Furthermore, our findings support that three other taxa belonging to the *O. vulgaris* group (*O. tetricus*, *O.* cf. *tetricus* and *O. sinensis*) and *O. insularis* are valid species. Our novel finding that analyses based on mtDNA does underestimate species diversity among octopuses shows that future phylogenetic studies require nuclear markers and/or be complimented with morphology-based evidence. Octopuses being exported globally under the name *O. vulgaris* are of extremely high market value and profile^33^. Aquaculture and captive growing of wild caught juveniles are receiving increasing profile and funding, particularly in China. Our data has, therefore, significant implications for the naming, marketing, value and documentation of commercially harvested octopuses in this species complex, and for the appropriate management and conservation of this highly valued fisheries resource.

## Methods

No live animals were experimented on during this study. Tissue samples were obtained via donation from existing museum or university collections, or taken from whole deceased animals that were purchased from fish markets.

### Data accessibility

Raw, unassembled RADseq data is hosted in DRYAD (doi:00.0000/dryad). Assembled RADseq data and the final alignments (RAD1 – 3) are also available (doi:00.0000/dryad).

### Sampling

Tissue samples of sub-mature and mature individuals belonging to the *O. vulgaris* species group (*O. vulgaris* s. s. (Mediterranean and north-east Atlantic), *O. vulgaris* Type II (southern Brazil) and *O. vulgaris* Type III (South Africa), *O. tetricus* (east Australia), *O.* cf. *tetricus* (west Australia) and *O. sinensis* (Asia)) and close relative, *O. insularis* were obtained from 16 localities around the world (Fig. 1, Table 2), and were stored in ~90% ethanol at −80°C until processing.

### Library preparation and sequencing

Genomic DNA was extracted from mantle or arm tissue (~1 mm^3^) using a DNeasy Blood and Tissue Kit (QIAGEN) according to the manufacturer’s instructions, except for the final elution which was repeated twice in a single aliquot of 57°C ‘low TE buffer’ to increase DNA yield. Where possible, skin was trimmed from tissue as a noticeable decrease in PCR efficiency was observed when skin was included with the tissue sample in the initial DNA extraction. Our ddRAD library was prepared using a modified version of the Peterson *et al.*^16^ protocol (available at michaelamor.com/protocols). Samples were randomly allocated to a position in a 96-well, round bottom PCR plate to minimise biases in sample preparation.

Genomic DNA was digested for 18 hours in 30 μL reactions composed of 20 units each of EcoRI-HF and ClaI restriction enzymes (New England Biolabs), 3 μL CutSmart buffer (New England Biolabs), 20 μL DNA solution (50 ng total DNA) and 4 μL H_2_O. The enzyme ClaI is known to be sensitive to CpG methylation, which may result in locus dropout. However, octopuses (1.2%) and other molluscs (2.0%) display relatively low methylation levels^51^. Furthermore, methylation has been shown to range from having an important effect in early development of O. vulgaris to having no effect in adulthood^52^. To minimise this impact, we sampled only sub-mature and mature individuals from each locality.

Barcodes/adapters were ligated to digested DNA fragments in 40 μL reactions composed of 30 μL DNA digestion solution, 1 μL T4 DNA ligase, 4 μL T4 DNA ligase buffer (New England Biolabs), 1 μL H_2_O, 2 μL common anti-sense adapter (2 μM stock concentration) and 2 μL sense adapter (2 μM stock concentration) which was unique to each sample. Ligation solutions were incubated at 16-18°C for 1 hour then 2 μL EDTA (0.5 M) was added to stop the reaction. Non-ligated adapters were removed using a 0.7× (DNA solution volume) AMPure XP (Agencourt) magnetic bead purification. The purified DNA pellets were suspended in 12 μL 40°C H_2_O.

PCR reactions were performed using 0.5 μM of each indexed sense and anti-sense primer and (10 μM stock concentration), 10 μL KAPA HiFi Real Time PCR master mix (KAPA Biosystems) and 9 μL size selected DNA solutions in a total reaction volume of 20 μL. PCR cycle conditions included a single initial denaturing step (98°C for 2 minutes) and 18 cycles of denaturing (98°C for 15 seconds), annealing (60°C for 30 seconds) and extension (72°C for 30 seconds). Amplified DNA solutions were purified using AMPure XP beads/PEG 6000 solution (0.7× DNA solution volume), quantified using a Qubit® 2.0 Fluorometer (Invitrogen) and DNA was pooled in equal proportions (15.4 ng per sample). The pooled library was electrophoresed on a 1.5% agarose gel, after which fragments between 300-350 base pairs (bp) were excised and purified using the Wizard® SV Gel and PCR Clean-Up System (Promega). The size selected library was diluted to 12 pM and sequencing was performed using a 600 cycle (paired-end) v3 MiSeq Reagent Kit on an Illumina MiSeq with 10% PhiX spiked into the run.

### Quality filtering and bioinformatics pipeline

Raw paired-end reads were merged using PEAR v0.9.4^53^. Merged and unmerged (read one only) reads were demultiplexed into individual sample read-sets based on their corresponding ligated inline barcode and indexed adapter using the ‘process_radtags’ feature of STACKS v1.2.7^54^. This step was also used to trim all reads to 250 bp as a noticeable decrease in read quality was recorded after 250 bp; any additional low quality reads were discarded (based on phred 30 quality score). The remaining good quality reads were then filtered to exclude microbial contamination using Kraken v0.10.4^55^ and unclassified, non-microbial reads were retained for phylogenetic analyses. Unmerged read two data were not included in the subsequent analyses as these reads did not meet the assumption that all reads are unlinked.

### Mapping to the mitochondrial genome

Demultiplexed filtered reads for each sample were mapped using Bowtie v2.2.2^56^ to the Japanese *O. vulgaris* mtDNA genome (Genbank accession NC006353). BAM files were generated using Samtools v0.1.19^57^. Mapped reads were imported into CLC genomics Workbench v7.0.4 (https://www.qiagenbioinformatics.com). A depth of five reads per individual surrounding restriction sites was required before consensus sequences were obtained manually. Multiple sequence alignment was performed on consensus sequences using the Muscle algorithm^58^. jModelTest v0.1.1^59^ was used to carry out statistical selection of best-fit models of nucleotide substitution of the alignment. The most appropriate substitution model (GTR+G) was selected based on ‘goodness of fit’ via the Akaike Information Criterion (AIC^60^).

Maximum likelihood analyses were performed using RAxML v8.0.19^61^. Strength of support for internal nodes of ML construction was measured using 1,000 rapid BS replicates. Bayesian Inference (BI) marginal PP were calculated using MrBayes v3.2.5^62^. Model parameter values were treated as unknown and were estimated. Random starting trees were used and analysis was run for 15 million generation sampling the Markov chain every 1,000 generations. After removing the initial 10% of samples, split frequencies within MrBayes as well as the program Tracer v1.6^63^ were used to ensure independent Markov chains had converged and reached stationarity.

### *A*ssembly of RAD loci

To assemble our loci we used a combined reference mapping and de novo assembly approach implemented in ipyrad v0.7.28^64^. Indexing and mapping to the available *O. bimaculoides*^35^ genome was performed using the Burrows-Wheeler Aligner (BWA)^65^ software package. To ensure our reads mapped to the *O. bimaculoides* genome, we allowed for up to 15% sequence difference at each locus. All unmapped reads were then assembled de novo at 85% similarity. Further sequence quality filtering was performed to convert base calls with a phred Q score of <33 (1 error per 1,000 bases) into Ns, whilst excluding reads with >5 Ns. A minimum of five reads per individual was required for clustering of putative loci, and those with fewer than five reads were excluded. Putative loci containing more than two alleles were excluded as potential paralogs. Any locus containing one or more heterozygous site across more than three samples was excluded. In cases where individuals were missing a given locus, gaps were replaced with Ns in the multiple sequence alignment output.

Three datasets were generated and analysed. For each comparison, RAD loci were filtered to ensure that for each locus, a genotype was present in at least 44-47% of the individuals. This level of missing data was chosen as it maximised the number of loci retained without influencing the phylogenetic position of individuals (details below). The first (RAD1, 298 loci, n = 34) contained all samples (*O. vulgaris* species complex, *O. tetricus*, *O.* cf. *tetricus* and *O. insularis*) and required 15 individuals per locus. The second data set (RAD2, 1,060 loci, n = 7/15) included only the two individuals with the highest number of reads per taxa (*O. insularis* included three individuals to include North Brazil and Ascension) and required seven individuals per locus. The final data set (RAD3, 604 loci, n = 26) included only individuals belonging to the *O. vulgaris* species complex, *O. tetricus* and *O.* cf. *tetricus* and *O. sinensis* and required 12 individuals required per locus. Maximum likelihood phylogenies were constructed using the GTRGAMMA model in RAxML v8.0.19^61^. Strength of support for internal nodes of ML construction was measured using 1,000 rapid BS replicates. We visualised the correlation between missing data and phylogenetic position to ensure that missing data did not influence the position of individuals throughout our phylogenies. This analysis was performed using the ‘plot.locus.dist’ function in the RADami^66^ package, which was implemented in R v3.1.0^67^. Finally, we screened our datasets for outlier loci using BayeScan v2.1^68^. The chain sample size was set to 50,000 with a 5,000 step burn-in pilot chain. All downstream analyses were performed on the entire dataset as no outliers were detected.

### Phylogenetic concordance among genome-wide loci

Support for sub-optimal topologies was investigated via the ‘partitioned RAD phylogenetic analysis’ approach^36^ using the RADami package^66^ in R v3.1.0^67^. To visualise the number of loci supporting the optimal tree relative to neighbouring sub-optimal trees generated from the RAD3 data-matrix, a candidate pool of 250 trees was generated via NNI for comparisons with the RAxML optimal tree (i.e. ‘best tree’). A set of unique trees for each locus was then generated by pruning the 251 trees to only those tips present in each locus. Site likelihoods for each locus-tree were calculated in RAxML v8.0.19^61^ under the GTR+G model. The likelihood of each tree was then plotted against the number of loci favouring and disfavouring each tree.

### Phylogenetic hypothesis testing

To assess confidence of our study’s optimal topology, relative to previously published mtDNA-based alternate topologies, the AU test^69^ was performed using the software package CONSEL^70^. Three topologies were investigated; the optimal topology generated by analysis of the O. vulgaris group via the RAD3 data set, an alternate hypothesis whereby *O. sinensis* was sister taxon to a clade containing *O. tetricus* and cf. *tetricus* (mtDNA; Amor *et al.*^30^), and a second alternate hypothesis whereby *O. vulgaris* Type III and *O. vulgaris* s. s. form a monophyletic clade (mtDNA; Guerra *et al.*^27^; Amor *et al.*^29^). Constrained topologies were constructed using the concatenated RAD3 data set based on 608 loci and 28 individuals. Site based likelihoods for each alternate tree were estimated using RAxML v8.0.19^61^ under the GTR+G evolutionary model. Significance values for the AU test and BS/PP values were calculated for each topology.

### Species tree estimation

Coalescent based species tree estimation on the RAD3 data-matrix was conducted following a quartet inference approach^71^ using the SVDQuartet command in PAUP v4.0a146 for Unix/Linux^72^ (Swofford 2003). The ML based analysis was performed using 100,000 randomly generated quartets and 1,000 BS replicates.

### Combined Bayesian inference and divergence time estimation

The consensus sequence for each species was obtained and aligned using Geneious v7.1.7^73^. Statistical selection of best-fit models of nucleotide substitution were undertaken using jModelTest v0.1.1^59^. The appropriate model (GTR+G) was chosen based on ‘goodness of fit’ via the AIC. Ultrametric topologies and divergence times among clades were co-estimated using BI analyses implemented in BEAST v2.3.1 and its associated LogCombiner and TreeAnnotator software^74^. All analyses were conducted using a log-normal relaxed clock^75^ under the Yule-speciation process^76^ with a normally distributed prior^77^. Topologies were estimated using random starting trees, gamma site models and log-normal relaxed clocks and were run for 750 million generations, sampling the Markov chain every 5,000. Three independent runs were performed for each evolutionary rate analysed and the resulting log and tree files were combined. Tracer v1.6^63^ was used to ensure Markov chains had reached stationarity, effective sampling size (ESS) was adequate (>1,000) and to determine the correct ‘burn-in’ for the analysis. The ‘maximum clade credibility tree’ was then obtained.

To estimate divergence times within the *O. vulgaris* species group we used the rate of 0.0036 neutral substitutions per million years in cephalopods^35^. We also estimated divergence times a substitution rates based on the distantly related molluscan bivalve family Arcidae Lamarck, 1809, which was similarly used in a recent study investigating the historical population expansion of the giant squid genus *Architeuthis*^78^. Three rates of evolution have been estimated from bivalves, each calculated using the nuclear marker histone 3: (1) 0.14-0.2%, (2) 0.09-0.1% and (3) 0.02-0.03% per million years^79^. As we were interested in estimating divergence times among a group of closely related species, we used the latter rate which was based on the calibration point at the split between the sub-genera *Anadara* s. s. and the sub-genus *Grandiarca*. Mean values at the mid-point of each rate range were calculated and the upper and lower range were set using a normally distributed prior. The two remaining bivalve rates of evolution were not used for our estimates as they were based on deeper phylogenetic calibration points; (1) the subfamilies Noetiinae and Striarcinae and (2) Anadarinae and the clade containing subgenera *Fugleria* and *Cucullaearca*.

## Acknowledgements

We thank CC Lu (Museum Victoria, Australia), Chih-Shin Chen and Chia-Hui Wang (National Taiwan Ocean University, Taiwan), Xiaodong Zheng and Yuanyuan Ma (Ocean University of China, China) and Eric Hochberg (Santa Barbara Museum of Natural History, USA) for project support. We also thank the following people for contributing tissue samples; Eduardo Almansa (Instituto Español de Oceanografía, Spain), Jorge Ramos Castillejos (University of Tasmania, Australia), Mandy Reid (Australian Museum, Australia) Ian Gleadall (Tohoku University, Japan), Xiaodong Zheng, Vladimir Laptikhovsky (Centre for Environment, Fisheries and Aquaculture Science, UK) and Warwick Sauer (Rhodes University, South Africa). A La Trobe University internal grant and a La Trobe Asia grant awarded to JMS funded experiments and travel. An Australian Biological Resources Study (ABRS) grant awarded to MDA assisted with the travel costs. A La Trobe University (School of Life Sciences) postgraduate publication grant awarded to MDA provided financial support during the writing of this manuscript. SRD was supported by an Illumina MiSeq grant. The Brazilian Institute of Environment and Renewable Natural Resources (IBAMA/ICMBio), the Brazilian National Research Council (CNPq559863/2008-0; 481491/2013-9) and Ciencias do Mar-CAPES (043/2013) are acknowledged for their logistic and financial support to TSL. MDA thanks Julian Finn and the staff of the Department of Marine Invertebrates at Museum Victoria, Australia for assistance in carrying out this project and Shannon Hedtke (La Trobe University, Australia) for assistance with phylogenetic hypothesis testing.

## Author contributions statement

MDA, MDN, AR, TSL and JMS obtained tissue samples. MDA, SRD and JMS conceived the experiments. MDA conducted experiments with assistance from SRD. Bioinformatics was performed by MDA, NEH and AJR. MDA wrote the manuscript. All authors reviewed the manuscript.

